# Coupling of autonomic and central events during sleep benefits declarative memory consolidation

**DOI:** 10.1101/195586

**Authors:** Mohsen Naji, Giri P. Krishnan, Elizabeth A McDevitt, Maxim Bazhenov, Sara C. Mednick

## Abstract

While anatomical pathways between forebrain cognitive and brainstem autonomic nervous centers are well defined, autonomic–central interactions during sleep and their contribution to waking performance are not understood. Here, we analyzed simultaneous central activity via electroencephalography (EEG) and autonomic heart beat-to-beat intervals (RR intervals) from electrocardiography (ECG) during wake and daytime sleep. We identified bursts of ECG activity that lasted 4-5 seconds and predominated in non-rapid-eye-movement sleep (NREM). Using event-based analysis of NREM sleep, we found an increase in delta (0.5-4Hz) and sigma (12-15Hz) power and an elevated density of slow oscillations (0.5-1Hz) about 5 secs prior to peak of the heart rate burst, as well as a surge in vagal activity, assessed by high-frequency (HF) component of RR intervals. Using regression framework, we show that these Autonomic/Central Events (ACE) positively predicted post-nap improvement in a declarative memory task after controlling for the effects of spindles and slow oscillations from sleep periods without ACE. No such relation was found between memory performance and a control nap. Additionally, NREM ACE negatively correlated with REM sleep and learning in a non-declarative memory task. These results provide the first evidence that coordinated autonomic and central events play a significant role in declarative memory consolidation.

## Introduction

It is now well established that specific electrophysiological central events during sleep support the transformation of recent experiences into long-term memories, i.e., memory consolidation. Another direction of research has demonstrated evidence for a critical role of autonomic activity during waking in memory and learning. We have recently shown that autonomic activity during sleep may also be related to consolidation. What is not known is whether the coupling of central and autonomic systems during wake or sleep plays a role in memory consolidation. Here, we identify a novel coupling between the autonomic and central nervous systems during sleep, but not wake, that predicts the outcome of the memory consolidation.

Research has consistently shown that a period of non-rapid eye movement (NREM) sleep yields greater memory retention of declarative memories (e.g., explicit, episodic memories) than a comparable period of REM sleep or waking activity ^1^. Several EEG features of NREM sleep have been linked with memory consolidation, with most studies focusing on spectral power in the slow wave activity (SWA, 0.5–4Hz) and sigma (12–15 Hz) bands, or specific events including hippocampal sharp wave-ripples (150–250 Hz)^2^, cortical slow oscillations (SO, 0.5–1 Hz), and thalamic sleep spindles (12–15Hz) ^3^. In fact, co-occurring SOs and sigma/spindles may be a key mechanism of memory consolidation during sleep ^4^.

In humans, increases in sigma power during the SO up-states has been shown following learning of a declarative memory task ^5^. Furthermore, pharmacologically increasing spindles with zolpidem resulted in greater coupling of spindles and SOs ^6^ and declarative memory improvements ^7^. In rodents, the replay of neural activity from encoding during sleep has been proposed to occur through the temporal coupling of thalamic spindles, hippocampal sharp wave ripples, and cortical slow oscillations ^8^. Thus, although a full mechanistic understanding of sleep-dependent memory consolidation is far from realized, research suggests that the coordination of brain rhythms from several cortical and subcortical regions may be critical.

A different line of research has demonstrated a significant contribution of the autonomic nervous system for memory consolidation during waking ^9^. These studies implicate the tenth cranial “vagus” nerve, which is the primary pathway of communication between the autonomic and central nervous systems. The vagus communicates information about peripheral excitation and arousal via projections to the brainstem, which then project to many memory-related brain areas including the amygdala complex, hippocampus and prefrontal cortex^10^. Descending projections from the PFC to autonomic/visceral sites of the hypothalamus and brainstem create a feedback loop allowing for bi-directional communication between central memory areas and peripheral sites ^11^. In male sprague-dawley rats, post-encoding vagotomy impairs memory ^12^. In humans, vagal nerve stimulation during verbal memory consolidation enhances recognition memory ^13^. Thus, the autonomic nervous system (ANS) appears to play a substantial role in waking memory consolidation.

We have recently shown that ANS activity during sleep also associated with memory consolidation ^14^. In this study, subjects were given a memory test before and after a daytime nap. Along with measuring central activity during sleep, we also measured ANS activity using the traditional approach of heart rate variability (HRV), defined as the variance between consecutive heartbeats averaged within each sleep stage, as well as during a pre-nap wake period. We found that vagally-mediated ANS activity during sleep 1) is associated with the consolidation of implicit and explicit information, and 2) is sleep stage specific.

Given the evidence of independent contributions of central and autonomic activity during sleep for memory consolidation, it is not known whether there is coupling between these systems and whether this coupling may support long-term memory formation. Prior work hints at a possible coordination between central features and autonomic activity. For example, auditory-evoked K-complexes were associated with increased heart rate ^15^ and have been shown to appear frequently 250 and 650 msec after the onset of the P wave in ECG ^16^. Furthermore, the QRS complex of ECG has been shown to modulate sleep spindle phases ^17^. There is also evidence that the heartbeat-evoked potentials in EEG reflect cardiac function ^18^. In addition, the high frequency component of heart rate variability, which reflects parasympathetic activity, has been shown to correlate with slow wave activity in the brain ^19^. In addition, volitional effort during wake correlates both with increases in hippocampal activity and heart rate^20^; and phase-locking between central hippocampal theta and autonomic R-waves has been shown in guinea pigs during wake, SWS and REM sleep ^21^. Together, these findings suggest that cardiac autonomic activity may be coupled with hippocampothalamocortical communication that has been shown to underlie memory consolidation during sleep. Despite these intriguing associations, very little is known about the coupling of central and autonomic activity and its functional consequences.

Here, we use a high-temporal precision time-series approach to examine coupling between central and autonomic nervous activity during wake and sleep and its impact on memory consolidation. Using this approach, we have identified novel cardiovascular events during NREM sleep, heart rate bursts, that are temporally coincident with increases in electrophysiological central events that have previously been shown to be critical for sleep-dependent memory consolidation. In addition, these Autonomic/Central Coupled Events (ACE) are directly following by a surge in vagal activity, assessed by high-frequency component of RR intervals. Using a regression framework, we assessed the contribution of ACE events versus non-ACE central and autonomic activity to declarative memory improvement following a nap, and show that ACE events predict performance improvement to a greater extent than by either activity alone. No such relation was found between memory performance and a control nap. We, thus, demonstrate a heretofore-unrecognized important role of coordinated autonomic and central activity during sleep that supports declarative memory consolidation.

## Results

We analyzed the RR (inter-beat interval) time-series from the ECG, and brain electrical activity from frontal and central EEG recording sites during a daytime nap in 45 young, healthy subjects (see Figure 1 for Study Timeline). First we assess HR using the traditional methods of analyzing HR in the frequency domain The total number of 3-minute epochs analyzed across subjects in Wake, Stage 2, SWS, and REM were 223, 415, 275, and 147, respectively. In the frequency domain, the low frequency (LF; 0.04– 0.15 Hz) component of RR is considered a reflection of sympathetic nervous activity ^22^; but this is not universally accepted ^23^, while the high frequency (HF; 0.15–0.4 Hz) component reflects parasympathetic (vagal) activity ^22^. Heart rate variability (HRV) analysis has revealed a decrease in LF and an increase in HF components in NREM during nighttime sleep ^24^ and naps ^25^. In agreement with this literature, we found that different sleep stages showed distinct properties in EEG and RR time-series frequency-domain. Peaks emerged in delta (0.5–4 Hz), theta (4–8 Hz), alpha (8–13 Hz), and sigma (12–15 Hz) bandwidth of EEG power spectrum (Figure 2b). In addition, the power spectrum of the RR time-series was modulated by sleep stages (Figure 2a).

**Figure 1.**
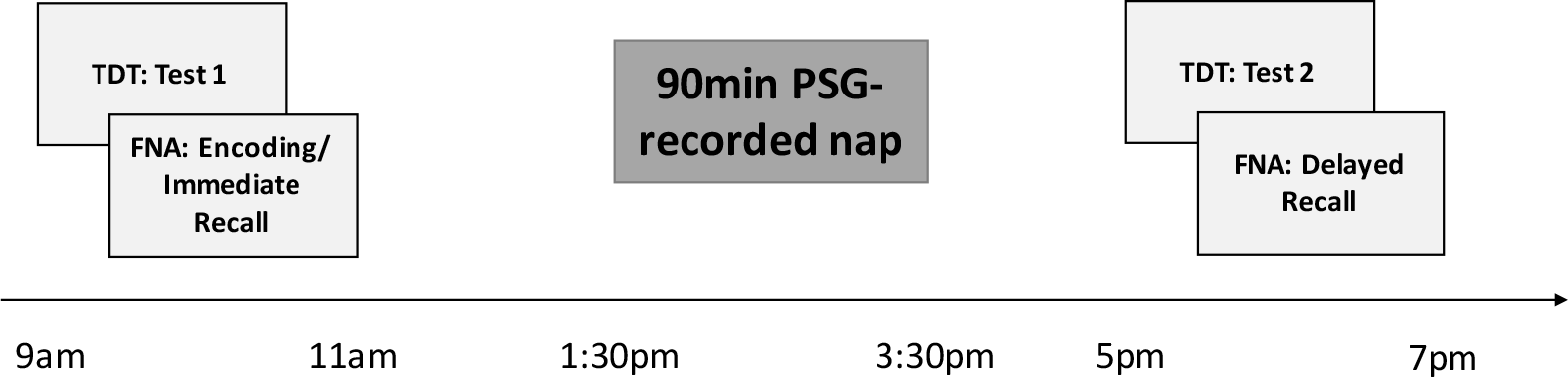
Study timeline for experimental nap. Subjects completed a declarative and non-declarative memory task. The order of tasks was counterbalanced across subjects. Before and after the daytime nap, declarative and procedural memory performances were tested.

We also employed a high-temporal precision time-series approach to the HR signal. In addition, after confirming the detected ECG R peaks by visual inspection, we intentionally analyzed all artifact-free RR intervals thereby retaining a larger amount of RR intervals than typical for HR analysis. Using this approach, we observed large bursts of HR (i.e., decreased RR intervals) over periods of 4-5 seconds (Figure 2c). Further, these HR bursts were visually noted to co-occur with events in the EEG (see boxes in Figure 2d–g). In the next section we provide a detailed analysis of these HR bursts and their coincidence with events in the EEG.

**Figure 2.**
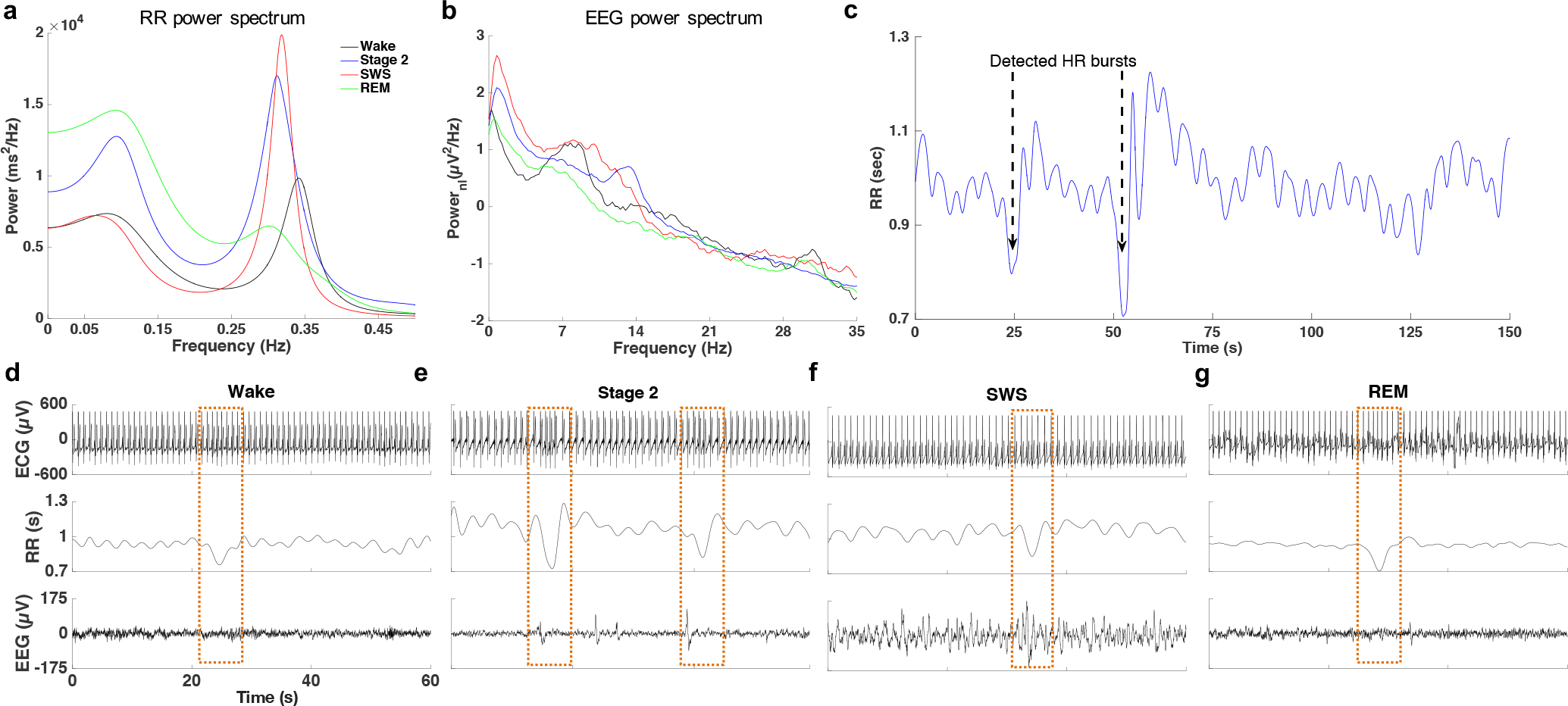
Characteristic properties of the EEG and RR time-series signals across wake and sleep stages (for one participant). a) RR time-series power spectrum during wake and sleep stages. b) EEG power spectrum (0–35 Hz) during wake and sleep stages. C) Detected HR bursts within a 150-sec bin during Stage 2. d-g) Simultaneous presentation of ECG, RR time-series, and raw EEG within 60-sec windows during wake, Stage 2, SWS, and REM, respectively. The boxes show the coincidence of HR bursts and EEG events during Stage 2 and SWS

## Temporal analysis of RR intervals reveals distinct, intermittent reductions in RR intervals

The HR bursts sporadically emerged in the time series of RR intervals (Figure 2c). The signature of these HR bursts was an increase in HR from baseline to the peak of the HR bursts (Wake: 21.62% (s.d.=11.05); Stage 2: 20.69% (s.d.=7.53); SWS: 14.49% (s.d.=6.02); REM: 21.40% (s.d.=8.85). The densities of the HR bursts for Wake, Stage 2, SWS, and REM were 0.71/min (s.d.=0.52), 1.04/min (s.d.=0.32), 1.11/min (s.d.=0.34), and 1.00/min (s.d.=0.38), respectively. Shapiro-Wilk and Chi-square tests on inter-event-intervals revealed that these events are generally aperiodic (Figure S1).

Following the visual observation of the co-occurrence of HR bursts and EEG events we assessed the degree to which the HR bursts correlated with changes in EEG activity (Figure 2c-d). We found that EEG delta amplitude during HR bursts was significantly higher than during non-bursting periods of the RR signal in both Stage 2 (t(84)=2.14, p=.035)) and SWS (t(70)=1.70, p=.046), but not Wake and REM. Traditional measures of HR in the frequency domain examine LF and HF power by collapsing across time. However, given our interest in moment-to-moment changes in ANS/CNS signals, we filtered the RR time-series by LF and HF frequency bands (RR_LF_ and RR_HF_, respectively), thereby maintaining the integrity of the time-series data. Similarly, we filtered the EEG time-series within the delta frequency to analyze the dynamics of the ANS/CNS interaction. We noted that during Stage 2 and SWS, bursts in delta amplitude co-occurred with large troughs in RR_LF_ (corresponding to HR bursts), which were followed by increases in RR_HF_ (see an example in Figure 3a). We further investigated this coincidence by examining the the phase/amplitude coupling (PAC) between delta and RR_LF_, where the slow frequency RR_LF_ provides the phase and the faster frequency delta provides the amplitude. During Stage 2 and SWS, the distribution of delta amplitude in the LF phase was non–uniform and peaked at a preferred phase (Figure 3b). Across participants, the average preferred phase was -70.1°(s.d.=26.7°) for Stage 2 and -79.4° (s.d.=67.6°) for SWS. This also indicates that the elevation in EEG delta activity preceded the peak of the HR bursts (which occurred at LF phase of 0º). Interestingly, when we compared the density of the HR bursts to the traditional FFT analysis of LF power (i.e., likely sympathetic activity) within Stage 2 and SWS, total LF power was not significantly associated with HR burst density in Stage 2 (r= -.04, p= .81; Figure 3c) or SWS (r= -.08, p= .64). We next set out to investigate the PAC between EEG and RR across a broader frequency range.

## EEG power is modulated by RR phase

We used the comodulogram method, utilizing normalized modulation index (nMI) as the PAC measure, to examine how the fast EEG signal was nested within the slower RR time-series signal across Wake, Stage 2, SWS and REM sleep. Conceptually, for each frequency pair in the comodulograms the modulation of EEG amplitude by RR phase was compared to amplitude-shuffled surrogate data. Figure 3e shows the phase of the RR time-series in frequencies below 0.2 Hz strongly modulated the amplitude in EEG in frequencies below 4 Hz, slow wave activity (SWA), during Stage 2. The same relation was apparent for SWS, but to a lesser extent (unpaired t-test between LF–SWA nMI of Stage 2 and SWS: (t(77)=3.21, p=.002) (Figure 3f). The LF–SWA modulation during Stage 2 and SWS was significantly higher than that of other RR/EEG frequency pairs (i.e., LF–theta, LF–alpha, LF–sigma, HF–SWA, HF–theta, HF–alpha, and HF–sigma; Stage 2: *F*_7,336_=5.83, p=.000002; SWS: *F*_7,280_=2.97, p=.005). Taken together, the time series analysis indicates that EEG SWA amplitude in Stage 2 and SWS was modulated by the LF component of the RR time-series (RR_LF_).

**Figure 3.**
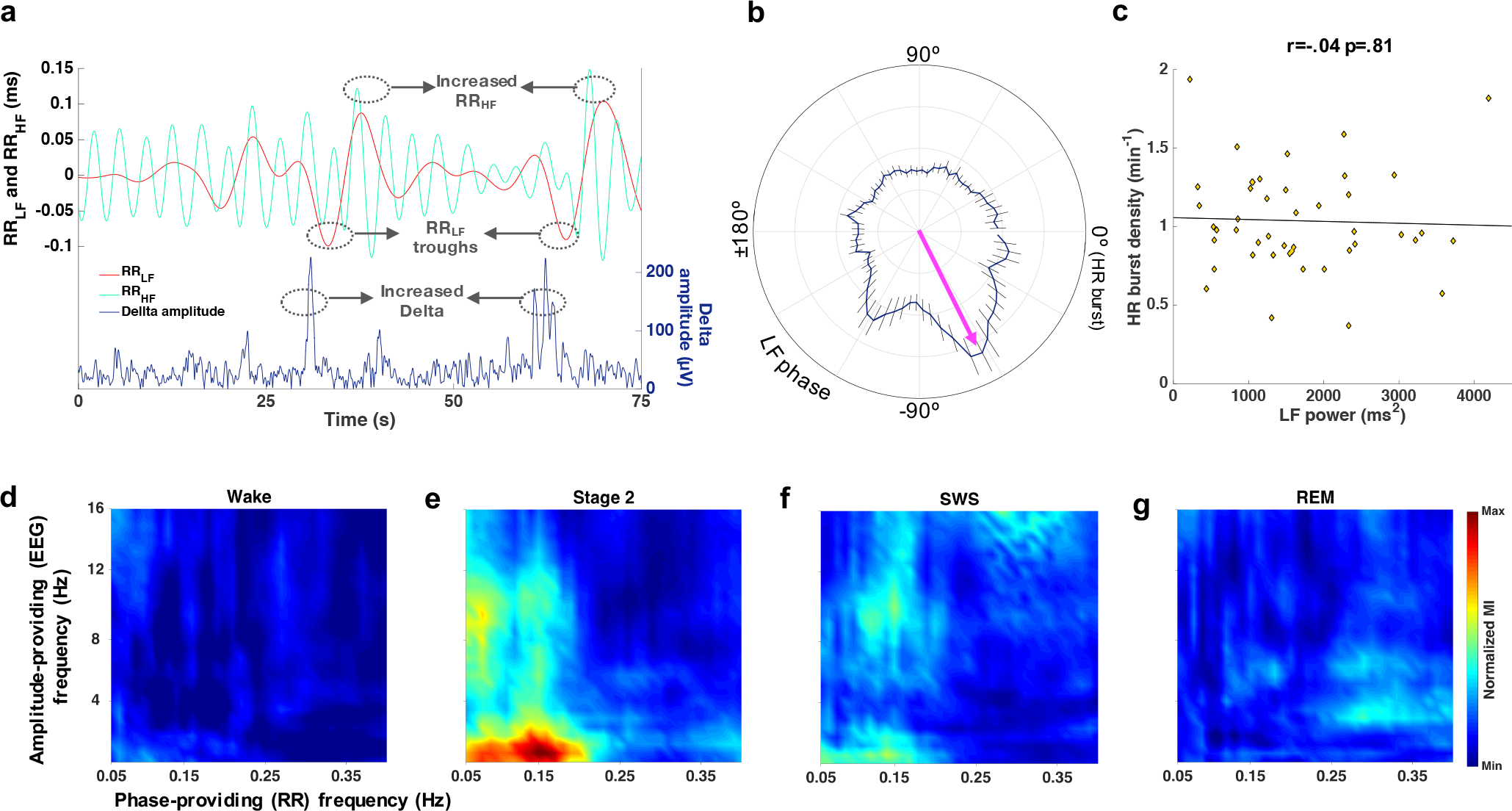
Temporal analysis of the RR intervals. a) A simultaneous presentation of delta amplitude and filtered components of RR time-series (i.e., LF and HF) showing the coincidence of large troughs in the LF component and elevated delta amplitude. b) The distribution of delta amplitude in LF phase is non-uniform and peaks at a preferred phase (LF troughs are assigned phase 0). c) The density of HR bursts does not significantly affect the LF power. d-g) The average EEG/ECG comodulograms, constructed from RR phase and EEG amplitude, across participants during wake, Stage 2, SWS, and REM, respectively. Error bars show standard error of the mean.

Though LF–SWA modulation was the strongest effect observed, the comodulogram also revealed other bands of EEG (<16 Hz) that were modulated by LF phase in Stage 2 compared with Wake and other sleep stages (LF-SWA: *F*_3,145_=10.44, p=.00001; LF-theta: *F*_3,145_=13.20, p=.004; LF-alpha: *F*_3,145_=17.58, p=.0005; LF-sigma: *F*_3,145_=13.93, p=.003). For REM sleep (Figure 3g), HF-modulated EEG theta activity was significantly higher than that of Wake (t(69)=4.85, p=.000007) and SWS (t(66)=2.06, p=.043) (*F*_2,103_=7.34, p=.002) but not Stage 2 (t(73)=0.45, p=.65). The other clusters visualized by the SWS comodulograms were not significantly different from Wake and sleep stages. In the following section, we will test the hypothesis that the above-mentioned modulation results from temporal coupling of autonomic and central events (ACE).

### Coordination between HR bursts and EEG

We investigated ACE coupling during wake and sleep stages by tracking fluctuations in the EEG in a 20-sec window from 10 second before to 10 second after the peak of the HR burst (Figure 4a–b). In addition to the rapid acceleration in HR, we also noted a slowing of HR after the burst or peak of the HR. We chose to use the peak of the HR burst as reference points because we found a larger number of HR bursts compared to HR troughs. The percentage of HR bursts which were followed by HR declines was 12.61±2.71 % during Wake, 20.30±2.14% during Stage 2, 13.30±2.65 % during SWS, and 27.02±4.00% during REM. 2). Therefore, the detection of the peaks was a more reliable metric of the HR bursts compared to detection of slower and smaller changes in HR. Furthermore, the magnitude of the HR slowing after the peak of HR was highly variable across HR bursts (see Figure S2).

As we were specifically interested in EEG activity related to memory consolidation, we narrowed our frequencies of interest to SWA (and SOs) and sigma activity (and spindles). EEG data were binned into 5-sec intervals within the 20-sec windows around the HR burst. Repeated measures ANOVAs indicated significant differences in SWA (Figure 4e) across the 5-sec bins and periods with no HR burst (baseline) during Stage 2 (*F*_4,168_=94.37, p<.00001) as well as during SWS (*F*_4,140_=21.59, p< .00001). Post hoc comparisons revealed the highest SWA occurred during the 5-sec bin prior the HR burst during Stage 2 (t(84)=8.94, p<.00001) as well as during SWS (t(70)=2.05, p=.044). Interestingly, the change in SWA in the 5-sec bin *prior* to the HR bursts was significantly correlated with the increase in HF power in the 5-sec bin *after* the HR burst (Figure 4c) during Stage 2 (r=.54, p=.002), likely reflecting a compensatory increase in vagal activity. This elevated increase in HF power after the HR burst was also present during other sleep stages (Figure 4c; Wake: t(44)=1.78, p=.082; SWS: t(70)=1.94, p=.056; REM: t(36)=2.12, p=.041, nonsignificant after FDR correction). Similar to SWA, density in SOs in the 5-sec bin prior to the HR burst was significantly increased in Stage 2 (Figure 4d, t(84)=8.83, p<.00001) as well as during SWS (t(70)=3.10, p=.007). Note that SWA without SO events (1-4Hz, 33.25±2.88 % of incidences) was also increased prior to the HR burst (t(82)=2.77, p=.003) during Stage 2. No significant correlation between HR bursts and SWA was found during Wake (*F*_3,112_=1.41, p=.24) or REM (*F*_3,116_=0.24, p=.87). To summarize, we found that ACE in Stage 2 and SWS, but not wake and REM sleep, were characterized by an increase in EEG SWA that occurred 5-sec before the peak of the HR burst and ended simultaneously with the HR bursts.

**Figure 4.**
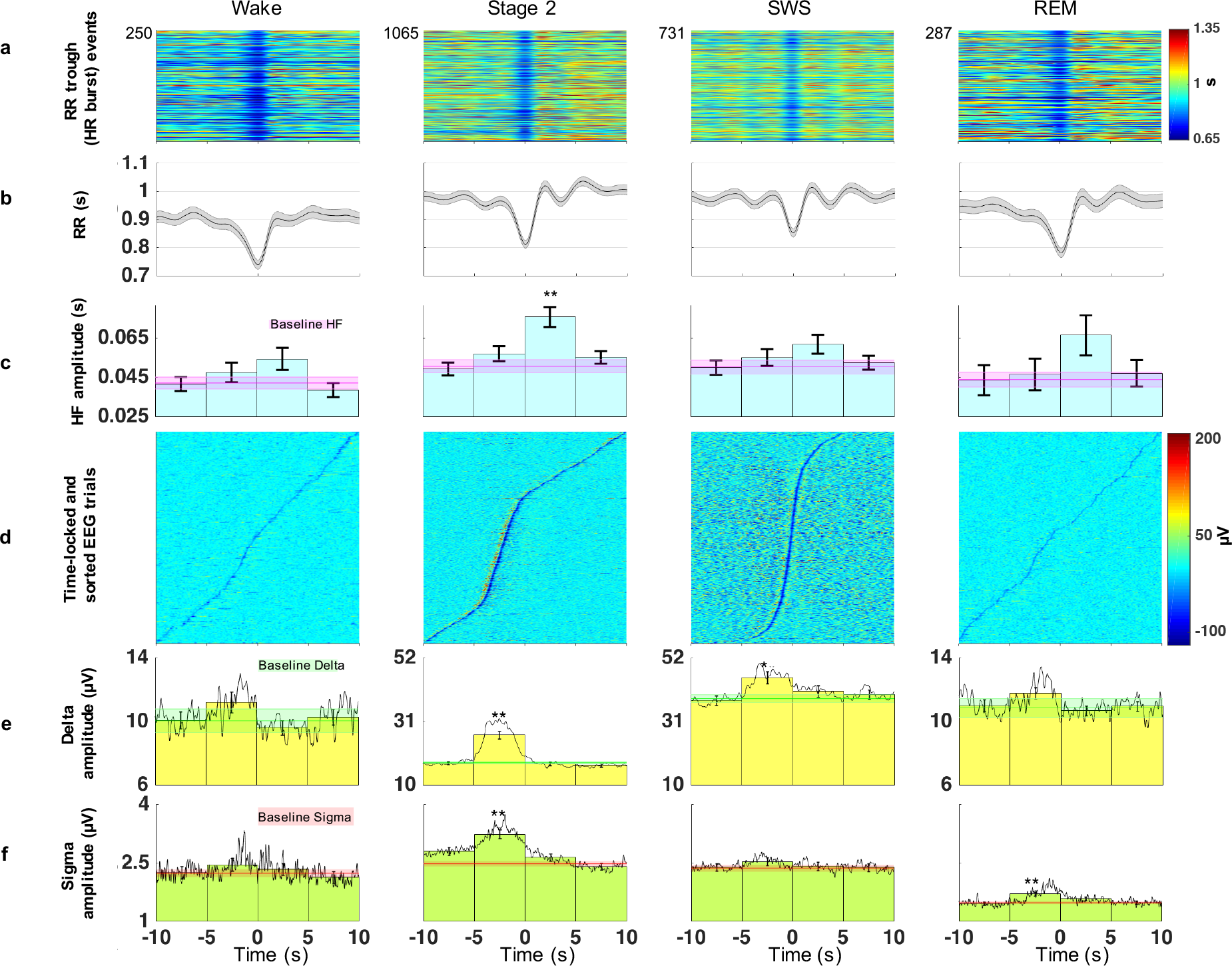
The event-related analysis of changes in ACE events. a) The HR burst events within a 20 s window for wake and different sleep stages. b) Grand average of the HR bursts. c) Average amplitude of HF component of the HR bursts in 5 s bins show a significant increase in the 5 s bin after the peak of the HR bursts. d) Event-locked EEG trials (Sorted based on the time difference between the HR burst at t=0 and the largest minimum of the EEG trials) show concentration of SOs prior the peak of HR bursts in NREM stages. e-f) average delta and sigma amplitude in 5 s bins (with the grand average of Delta amplitude on top of them) around the HR bursts, respectively. Asterisks in (c), (e), and (f) show the significant differences after FDR correction (*p<.05 and **p<.001) between an amplitude in a bin and the average amplitude in periods with no HR burst (baseline). Error bars show standard error of the mean.

Repeated measures ANOVAs also indicated significant differences in sigma activity (Figure 4f) across the 5-sec bins and periods with no HR burst during Stage 2 (*F*_4,168_=56.87, p<.00001). The highest sigma power occurred during the 5-sec bin prior the HR burst (t(84)=4.71, p=.00001). The early change in sigma power (10 secs prior the peak of the HR bursts) was not significant after FDR correction (t(84)=1.90, p=.061; Figure 4f). Similarly, repeated measure ANOVA indicated significant differences in spindle density across the 5-sec bins and periods with no HR burst (*F*_4,168_=2.93, p=.022, Figure S3). Post hoc paired-samples t-tests between baseline spindle density and the spindle density during the 5-sec bins revealed a significant density difference during the 5-sec bin prior the HR burst peak (t(84)=3.679, p=.0004) and a nonsignificant significant difference during the 5-sec bin started 10 sec before the HR burst peak (t(84)=1.871, p=.065). No significant modulation in sigma power was found during SWS (t(70)=0.17, p=.86).

In addition to SWA and sigma activity, the coupling of SO and spindles as a function of the HR was investigated. For each subject, the average SO/spindle modulation index ^26^ was calculated for two groups of SOs: 1) SOs occurred during the 20-sec windows around HR bursts and 2) SOs during periods with no HR burst. Paired t-test showed no difference in MI for the two SO groups (t(87)=0.177, p=.860). *That is, ACE-related sleep activity did not have any measurable impact on the temporal coupling of SO and spindles*.

In summary, Sigma and SWA power in Stage 2 and SWA in SWS increased from baseline (periods with no HR burst) to a maximum level prior to the peak of the HR bursts, and returned to baseline post-burst. Although we focused on slow wave and sigma frequencies here, we conducted exploratory analyses on theta and beta activity, which are presented in the Supplementary Materials. These data are consistent with the hypothesis that cortical EEG activity precedes and perhaps catalyzes these sudden and short-lived surges in HR in NREM sleep. We next investigated the functional impact of ACE events on memory consolidation during sleep.

## Correlations with cognitive tasks

In this section, we investigated the contribution of ACE coupling to post-nap memory consolidation. For this purpose, we computed the change scores for both performance and EEG characteristics, which were calculated as average values 5-secs prior to the peak of the HR bursts subtracted from baseline (ACE difference scores). We focused on Stage 2 and SWS, as ACE was not significantly modulated during wake or REM. The following characteristics were analyzed: density of SOs and magnitude of SWA and sigma power in Stage 2 and SWS, and a linear composite of SWA and sigma power changes (i.e., a simple sum of z-scores of SWA and sigma power changes) in Stage 2 (Figure 5). We conducted exploratory analyses on the application of modulation index between HR bursts and EEG activities, which are presented in the Supplementary Materials. (see Figure S5). Two memory tasks were considered for this study: declarative memory for face-name associations and non-declarative perceptual learning on a texture discrimination task. We calculated difference scores between pre-nap and post-nap memory performance for total (first and last name) recall and perceptual learning thresholds.

### Declarative Memory

Recall difference scores were positively correlated with ACE difference scores in density of SOs in Stage 2 (r= .47, p= .002, significant after FDR correction; Figure 5a) and SWS (r= .47, p= .004, significant after FDR correction; Figure 5d), as well as SWA (r= .32, p= .039, marginally significant after FDR correction; Figure 5b), sigma power (r= .3, p=.05; Figure 5c), and the linear composite of SWA and sigma power in Stage 2 (r= .38, p=.012, significant after FDR correction; Figure 5f). The correlation between changes in performance and SWA in SWS was not significant (r= .11, p= .54; Figure 5e). Additionally, the ACE difference score for HF power in Stage 2 sleep (5-sec bin *after* the peak of the HR burst) was significantly correlated with recall improvement (r=.32, p=.03, marginally significant after FDR correction).

**Figure 5.**
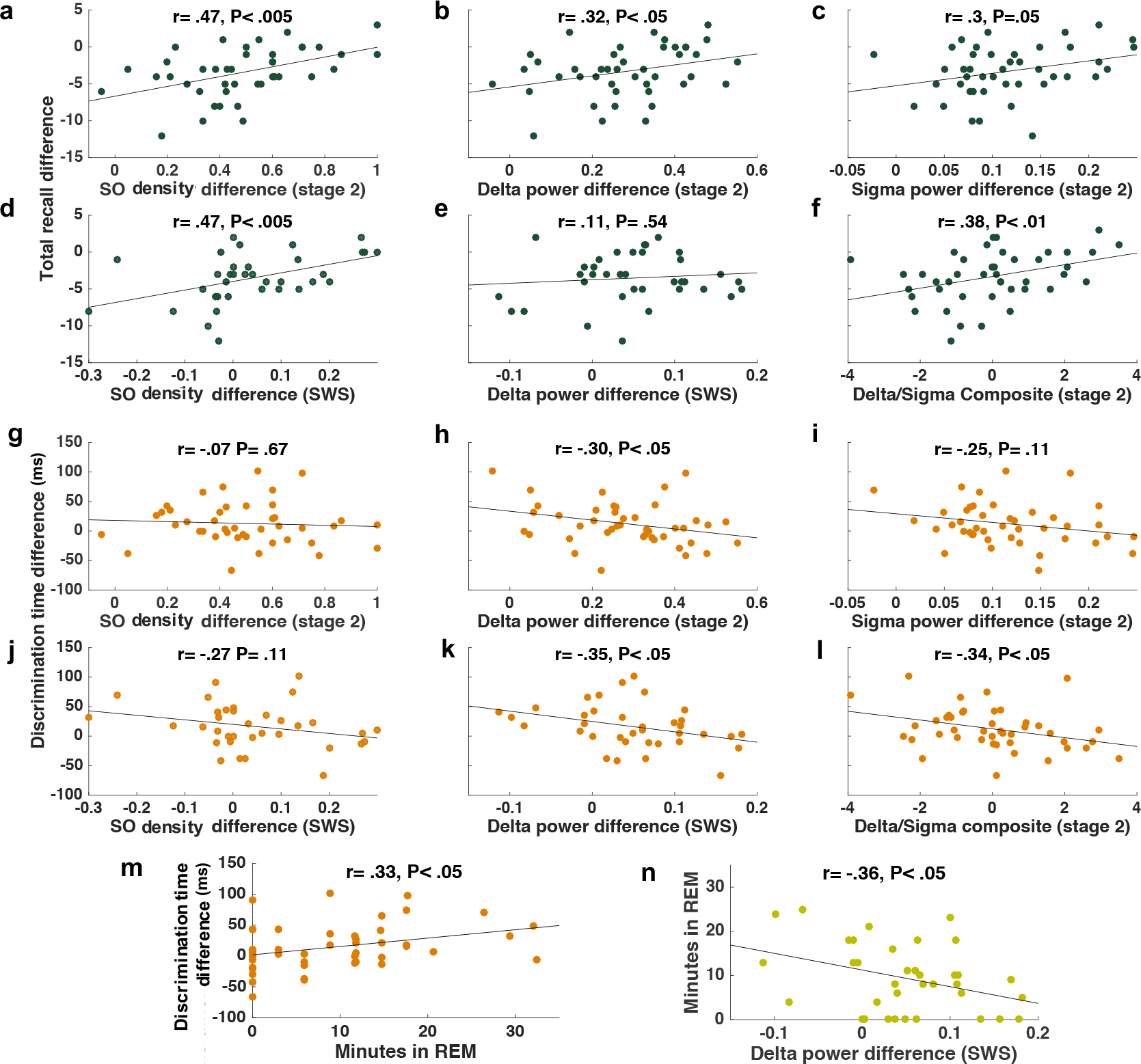
Impact of ACE on memory consolidation. a-f) Scatter plots for relationships between the recall improvement in the declarative memory (face-name task) and ACE difference scores of SO density, delta power, and sigma power during Stage 2 (n=42) and SWS (n=36). Note, that performance was positively correlated with increase in ACE difference scores. g-l) Scatter plots for relationships between improvement in the perceptual learning (texture discrimination task) and ACE difference scores of SO, delta power and sigma power during Stage 2 and SWS. Note, negative correlation in all cases. m-n) Scatter plots for relationships between minutes in REM sleep and texture discrimination task learning and ACE difference scores of delta power in SWS, respectively.

To assess the relative importance for memory performance of independent autonomic and central events as well as their coupling, we utilized a hierarchical, linear regression framework. For Stage 2 and SWS, two linear regression models were calculated to predict recall difference. In Model 1, SO density, spindle density, burst density, and baseline HF power in each sleep stage were the independent variables. In Model 2, we added the ACE difference scores for SO density and spindle density *before* and HF power *after* the HR bursts. The regression results for Stage 2 and SWS variables are tabulated in Table 1 and Table 2, respectively. Stage 2 results showed that Model 1 was not significant (*F*_4,37_=0.08, p= .99; *adj* R^2^ = -.10), whereas Model 2 significantly predicted performance (*F*_7,34_=2.77, p= .022; *adj* R^2^ = .23) with both ACE SO density change and HF amplitude change as significant predictors. Comparing Model 1 and 2, we found that Model 2 explained significantly more of the variance in recall than Model 1 (change in *adj* R^2^=.33, *F*_3,34_=6.30, p=.002). For SWS, Model 1 was not significant (*F*_4,30_=.36, p=.84; *adj* R^2^ = -.08), but adding the ACE measures in Model 2 elevated the model to a marginal significance level (*F*_7,27_=1.53, p=.20; adj R^2^ = .10), with SO density change the only significant predictor. Again, Model 2 accounted for significantly more of the variance in recall improvement than Model 1 (change in adj R^2^=.21, F_3,27_=3.01, p=.048). In summary, while the baseline HF power, SO, spindle, and HR burst densities did not independently predict recall difference, ACE events predicted up to 23% of the variance in performance improvement on this declarative memory task.

**Table 1:** Regression models from Stage 2 variables to predict FNA total recall difference

| Variables | Model 1 |  |  |  | Model 2 |  |  |  |
| --- | --- | --- | --- | --- | --- | --- | --- | --- |
| | B | SE | t | $\beta$ | B | SE | t | $\beta$ |
| HR burst density | 0.39 | 1.84 | 0.21 | 0.04 | -1.69 | 1.66 | -1.02 | -0.15 |
| SO density | -0.13 | 0.40 | -0.31 | -0.06 | -0.12 | 0.35 | -0.35 | -0.05 |
| Spindle density | -0.16 | 0.90 | -0.22 |  | 0.61 | 0.81 | 0.76 | 0.11 |
| Baseline HF amplitude | -5.79 | 26.32 | -0.22 | -0.04 | 19.98 | 23.53 | 0.85 | 0.13 |
| SO density change score |  |  |  |  | **6.27** | **2.26** | **2.77** | **0.44\*\*** |
| Spindle density change score |  |  |  |  | 2.17 | 0.81 | 1.09 | 0.17 |
| HF amplitude change score |  |  |  |  | **8.56** | **3.46** | **2.47** | **0.38\*** |
| (Constant) | -2.83 | 3.32 | -0.85 |  | -8.94 | 3.22 | -2.77 |  |
| F |  |  | 0.08 |  |  |  | **2.77\*** |  |
| R<sup>2</sup> |  |  | 0.01 |  |  |  | 0.36 |  |
| Adjusted R<sup>2</sup> |  |  | -0.10 |  |  |  | 0.23 |  |
| Change in Adjusted R<sup>2</sup> |  |  |  |  |  |  | **0.33\*\*\*** |  |

\*p < .05, \*\*p<.01, \*\*\*p<.001

**Table 2:** Regression models from SWS variables to predict FNA total recall difference

| Variables | Model 1 |  |  |  | Model 2 |  |  |  |
| --- | --- | --- | --- | --- | --- | --- | --- | --- |
| | B | SE | t | $\beta$ | B | SE | t | $\beta$ |
| HR burst density | -0.43 | 0.55 | -0.78 | -0.15 | -0.83 | 1.69 | -0.49 | -0.08 |
| SO density | -0.01 | 0.02 | -0.53 | -0.10 | 0.03 | 0.08 | 0.42 | 0.07 |
| Spindle density | -0.13 | 0.21 | -0.62 | -0.11 | -0.04 | 0.64 | -0.06 | -0.01 |
| Baseline HF amplitude | 2.02 | 8.49 | 0.24 | 0.04 | 21.97 | 27.09 | 0.81 | 0.14 |
| SO density change score |  |  |  |  | **11.85** | **4.27** | **2.77** | **0.48\*\*** |
| Spindle density change score |  |  |  |  | 0.09 | 1.37 | 0.07 | 0.01 |
| HF amplitude change score |  |  |  |  | -9.11 | 7.38 | -1.23 | -0.21 |
| (Constant) | -2.83 | 3.32 | -0.85 |  | -8.94 | 3.22 | -2.77 |  |
| F |  |  | 0.36 |  |  |  | 1.53 |  |
| R<sup>2</sup> |  |  | 0.05 |  |  |  | 0.28 |  |
| Adjusted R<sup>2</sup> |  |  | -0.08 |  |  |  | 0.10 |  |
| Change in Adjusted R<sup>2</sup> |  |  |  |  |  |  | **0.18\*\*\*** |  |

\*p < .05, \*\*p<.01, \*\*\*p<.001

### Perceptual Learning

We found perceptual learning was negatively related to the ACE SWA increase in Stage 2 (r=- .30, p= .048; Figure 5h) and SWS (r= -.35, p= .035; Figure 5k), as well as with the linear composite of SWA and sigma power changes in Stage 2 (r= -.34, p=.025; Figure 5l). In addition, non-significant negative correlations were apparent between perceptual learning and ACE SOs increases in Stage 2 (r= - .07, p= .67; Figure 5g) and SWS (r= - .27, p= .11; Figure 5j) and Stage 2 sigma power (r= -.25, p=.11; Figure 5i). Thus, in contrast with the positive association between ACE-mediated increases in NREM sleep events and declarative memory, perceptual learning was negatively associated with these ACE events. Given that prior studies have demonstrated the critical importance of both NREM and REM sleep for perceptual learning ^27^, the trade-off between ACENREM features supporting declarative memory and REM features supporting perceptual learning may not be surprising. Indeed, Figure 5m shows a similar significant positive correlation between REM sleep and perceptual learning (r=.37, p=.013, marginally significant after FDR correction). We also showed a reciprocally negative correlation between ACE SWA difference score and minutes in REM (r= -.36, p=.029; Figure 5n). These negative associations with REM and REM-dependent learning may be related to the experimental conditions of the nap, in which subjects have a two-hour nap opportunity. Such restrictions on sleep provide boundaries on the total sleep time and may thus create a trade-off between NREM and REM sleep.

In summary, we find that sleep features associated with consolidation of hippocampal-dependent memories (SO events, SWA, and sigma power) are specifically boosted during HR bursts and that these ACE increases may be an important contributor in hippocampal-dependent memory consolidation. However, these benefits to declarative memory during a nap may come at the expense of REM sleep and REM-dependent perceptual learning.

### Control group

A subset of subjects (n=22) were given a control nap one week after the experimental nap. On both experimental and control days, subjects were tested on cognitive tasks in the morning and evening and had a nap between test sessions, however, different cognitive tasks were tested on these days. This is an appropriate within-subjects control because the subjects had the same magnitude of cognitive burden, but with different information to encode, which allows us to test the specificity of the relationship between changes in the autonomic/central events during sleep and the specific memories learned. Underscoring this point, we found similar levels of HR burst density (0.97/min) and changes in HF power (control: 20.07%, experimental: 18.61%), delta power (control: 17.65%, experimental: 21.37%), and sigma power (control: 7.64%, experimental: 8.63%) in the experimental and control naps during Stage 2 and SWS.

Next, we confirmed that these subjects had the same magnitude of effect size for the association between nap ACEs and memory on the experimental day as the entire sample. We re-ran the regression models from Table 1, in which Model 1 compared non-ACE predictors to declarative memory, and Model 2 added the ACE predictors. Indeed, a model with baseline variables did not significantly predict the declarative learning (*adj* R^2^ = -.23, p=0.95), whilst adding ACE difference scores in Model 2 significantly improved the prediction (*adj* R^2^ = .47, p=0.04).

Finally, in order to confirm the specificity of the encoded material on the experimental day, correlations between control nap ACE difference scores and memory changes on the experimental day were tested. Here, evidence of no relation between the control nap ACE features and experimental memory performance would indicate a high degree of specificity to the encoded material on the experimental day. For the declarative memory task, no significant correlation was found with ACE SO density difference score in control nap Stage 2 (r=-0.04, p=0.85) and SWS (r=-0.14, p=0.61), as well as with the SWA changes in Stage 2 (r=0.26, p=0.24) and SWS (r=-0.35, p=0.16), and sigma power changes in Stage 2 (r=0.11, p=0.63) and SWS (r=0.14, p=0.60). Unlike experimental day, there was no correlation between declarative learning and HF power change in Stage 2 (r=0.15, p=0.49). Similarly, no significant correlation was found between perceptual learning and ACE SO density increase in Stage 2 (r=0.05, p=0.82) and SWS (r=-0.18, p=0.49), as well as with the SWA changes in Stage 2 (r=0.02, p=0.93) and SWS (r=0.07, p=0.78), and sigma power changes in Stage 2 (r=0.12, p=0.59) and SWS (r=0.08, p=0.75). In summary, ACE changes in the control nap were not associated with memory consolidation of information encoded on the experimental day.

## Discussion

Here, we have identified for the first time an autonomic cardiac event during NREM sleep that is temporally-coupled with a significant boost in central oscillations associated with systems consolidation. Specifically, we show 1) increases in the EEG amplitude in SWA and sigma bands directly preceded these large-amplitude HR bursts, 2) a surge in vagal activity (measured in the HF component of heart rate) directly following the HR burst, 3) that the uptick in autonomic/central events (ACE) (SO, delta, sigma) and vagal activity were positively associated with declarative memory improvement and negatively associated with non-declarative memory, and 4) ACE changes in the experimental nap were only associated with memory consolidation of information encoded on the experimental day, and not merely correlational. Together these results present compelling evidence of a coupling between signals from the brain and heart that have a critical and specific impact on memory consolidation during sleep.

Traditional calculations of EEG/ECG signals in the frequency domain using 3–5 min bins of uninterrupted nocturnal sleep have found results that differ from the present outcomes. One such study reported no association between delta power and normalized HF power at peak of SWA ^28^. Also different from the present result, cardiac changes have been found to *precede* EEG changes by several minutes ^29^. Here, unlike typical HRV analyses that discard large changes in HR, we intentionally analyzed all artifact-free RR intervals, which allowed for the discovery of these HR bursts. In addition, we used a time-domain analysis within short 20 sec windows around the HR bursts to examine fluctuations in the EEG/ECG signals, which allowed for more fine-grained assessment of event-related changes in both signals. Similarly, using an event-related analysis in near-infrared spectroscopy, Mensen and colleagues ^30^ reported an oscillating blood flow signal at HR frequency that was time-locked to the onset of slow waves. This study posited that the arterial pulsation evokes a down-state, or that a third generator regulating HR and slow waves may be involved. More in line with our findings, another event-based analysis revealed an increase in HR followed by a deceleration *after* spontaneous and tone-induced K-complexes ^15^.

By focusing on fine resolution of HR events for our event-based analysis, we found distinct bursts in HR during wake and sleep. HR bursts of similar duration have been identified in REM sleep in cats, with the average incidence rate of 1 burst per 6.1 min of REM sleep. These HR bursts were accompanied by ponto-geniculo-occipital waves and theta activity ^31^, which have been associated with memory consolidation in rats ^32^. Our event-based analyses showed SWA modulation in both Stage 2 and SWS, and sigma modulation in Stage 2 by the LF component of the RR time-series. Our results suggest that the mechanism of SWA modulation was an increase in density of SOs, and power in SWA, and sigma activity ~5 sec prior to the peak of the HR bursts that returned to baseline directly after the HR bursts. In addition, there was no significant SWA modulation in wake and REM, which is not surprising due to low SWA activity in these stages. The characteristics of the SWA and sigma modulation in NREM sleep resemble the cyclic alternating pattern (CAP) ^33^ in NREM sleep which is characterized by repeated sequences of transient events and which clearly break away from the ongoing background rhythm recurring at intervals up to 1 min long. However, the average duration of SWA reactivation in CAP was reported as 12.66 s which is longer than our ACE observations. Lecci et al ^34^ found periodic sigma activity (but not SWA) that was more prevalent in Stage 2 sleep than SWS and associated with memory consolidation in a declarative task (human) as well as hippocampal ripple activity (in mice). The present results, on the other hand, identified a heretofore undescribed burst in HR activity, that correlated with prominent, aperiodic, ACE-related increases in SWA and sigma, as well as parasympathetic activity, that predicted gains in declarative memory during Stage 2 and SWS. Differences between findings may be due to the focus of the Lecci paper on periodic changes in EEG activity, as opposed to using ECG as the reference point for analysis. Notwithstanding these differences in approach and outcomes, both studies point to an important link between memory consolidation and autonomic/central events during sleep.

The present data also showed no significant correlation between HR bursts and traditional frequency based analysis of total LF power. This lack of relation may suggest a dissociation between mechanisms underlying LF-related sympathetic activity and HR bursts. In contrast, the increase in HF power directly following the HR bursts may instead implicate vagal inhibition ^35^. Although specific mechanisms of ACE events are not known, we speculate that overlapping neural pathways between brain and heart centers may be an appropriate place to begin, and further investigation is needed to tease apart these mechanisms

Several candidate brainstem and cortical regions may be involved in the interaction between HR and sleep oscillations. One possibility, is that the nucleus of the solitary tract (NTS) and the rostral ventrolateral medulla (RVLM) mediate this interaction, since they act as one of the main bridges between CNS and cardiovascular systems ^36^. NTS is one of the critical components of the central autonomic network with afferent and efferent connections to the cardiovascular system ^37^ and it receives projections from many cortical regions ^38^. Further, NTS, through its projections to RVLMs, influences activity in the locus coeruleus ^39^ and indirectly influences the basal forebrain. Indeed, increased NTS activity is associated with reduced HR, and NTS stimulation has been shown to augment EEG theta and beta power during wake in cats ^40^ potentially through the input to RVLM. The HR burst may be a consequence of increases in spindle and slow oscillations through wide spread cortical projection to NTS ^38^. On the other hand, one potential mechanism for SO reduction post-HR burst may be via NTS projections to basal forebrain and locus coeruleus. Stimulation of these areas by the NTS increases acetylcholine and norepinephrine release in cortex, which is known to reduce slow oscillations ^41^. Taken together, this suggests that the increase of SO prior to HR burst could mediate an increase of HR through combination of sympathetic and parasympathetic pathways and that the reduction of SO following HR burst may arise from increased activity of NTS through parasympathetic output via the vagal nerve. However, several other pathways are known to be involved in the interaction between neocortex and heart and have be examined to make definite conclusions.

A large prior literature suggests a critical role of the ANS during wake for memory consolidation ^9^. Studies have found that direct modification of peripheral hormonal activity following acquisition can enhance or impair the memory storage of new information ^42^ via vagal afferent nerve fibers, which communicate information about ANS excitation via projections to the brainstem, which then project to memory-related areas including the hippocampus, amygdala complex, and prefrontal cortex ^10^. Bidirectional projections from the prefrontal cortex to the hypothalamus and brainstem create a feedback loop for communication between peripheral sites and central memory areas ^11^. Lesions of the vagus nerve impair memory ^12^, whereas pairing vagal nerve stimulation with auditory stimuli reorganizes neural circuits ^43^, strengthens neural response to speech sounds in the auditory cortex ^44^, and enhances extinction learning of fear memories in rodents ^45^. In humans, vagal nerve stimulation enhances consolidation of verbal memory ^13^ and working memory ^46^. Recently, Whitehurst and colleagues used traditional methods to detect total sympathetic and parasympathetic power of the HRV signal during sleep and showed a correlation between parasympathetic activity during REM sleep (but not SWS) and implicit learning ^14^. In contrast the current study used a fine-grained temporal analysis of EEG/ECG signals to reveal how the coupling of ANS/CNS Events (ACE) significantly improved the prediction of declarative memory, but not non-declarative memory, over and above total power of HR components and sleep events alone. Importantly, these effects are specific to the encoded memories directly preceding the nap. Follow-up experiments require interventions to better understand the mechanism promoting these effects, as well as probing different clinical populations that may have dampened autonomic tone during sleep, including older adults and sleep apnea. Taken together, these findings implicate a significant role of ANS modulation of plasticity and memory consolidation during sleep that is likely mediated by vagal afferents to the cortex via NTS brainstem.

## Methods

### A. Participants

Data reported here come from the first visit of a larger, mini-longitudinal study that included up to 7 visits per participant. Data from this study have been reported elsewhere ^14, 25 47^. Fifty-five (30 females) healthy, non-smoking adults between the ages of 18 and 35 with no personal history of sleep disorders, neurological, psychological, or other chronic illness gave informed consent to participate in the experimental nap protocol explained below. Twenty-two subjects also participated in a one-day control nap study. All experimental procedures were approved by the Human Research Review Board at the University of California, Riverside and were in accordance with federal (NIH) guidelines and regulations. Participants were thoroughly screened prior to participation in the study. The Epworth Sleepiness Scale (ESS ^48^) and the reduced Morningness-Eveningness questionnaire (rMEQ ^49^) were used to exclude potential participants with excessive daytime sleepiness (ESS scores >10) or extreme chronotypes (rMEQ <8 or >21). Additionally, potential participants were interviewed about their medical history and quality and quantity of their sleep, including questions from the DSM-IV used to diagnose sleep-wake disorders. Participants included in the study had a regular sleep-wake schedule (reporting a habitual time in bed of about 7–9 h per night), and no presence or history of sleep, psychiatric, neurological, or cardiovascular disorders determined during screening. Participants received monetary compensation and/or course credit for participating in the study.

### B. Data acquisition and pre-processing

#### - Study Procedure

Participants wore actigraphs to monitor sleep-wake activity for one week prior to the experiment to ensure participants were not sleep-deprived and spent at least 6.5 hours in bed the night prior to their visit. On both the experimental and control nap days, subjects arrived at the UC Riverside Sleep and Cognition lab at 9AM for Test Session 1. On the experimental day, cognitive tasks included a declarative and non-declarative memory task (see below), as well as a creativity measure (reported elsewhere). On the control day, subjects were tested in a spatial navigation task and verbal memory task. The order of tasks was counterbalanced across subjects. In total, the declarative memory task took about 40 minutes to complete, and the non-declarative task took about 20 minutes to complete (the creativity measure took about 50 minutes, and all testing was done by approximately 11AM). Between completing the tasks and starting the nap, participants were allowed to leave the lab and go about their normal activities, except avoiding caffeine and napping. At 12:30PM, electrodes were attached for polysomnography (PSG) recording. At 1:30PM, subjects took a PSG-recorded nap. They were given up to 2 hours time-in-bed to obtain up to 90 min total sleep time. Sleep was monitored online by a trained sleep technician. Nap sessions were ended if the participant spent more than 30 consecutive min awake. Naps were completed at approximately 3-3:30PM, electrodes were removed, and participants were given a break where they could leave the lab. At 5PM, subjects returned to the lab for Test Session 2.

#### - Sleep recording

Polysomnography (PSG) data including electroencephalogram (EEG), electrocardiogram (ECG), chin electromyogram (EMG), and electrooculogram (EOG) were collected using Astro-Med Grass Heritage Model 15 amplifiers with Grass GAMMA software. Scalp EEG and EOG electrodes were referenced to unlinked contralateral mastoids (F3/A2, F4/A1, C3/A2, C4/A1, P3/A2, P4/A1, O1/A2, O2/A1, LOC/A2, ROC/A1) and two submental EMG electrodes were attached under the chin and referenced to each other. ECG was recorded by using a modified Lead II Einthoven configuration. All data were digitized at 256 Hz.

#### - Sleep scoring

Raw data were visually scored in 30-sec epochs according to Rechtshaffen and Kales ^50^. Five sleep stages (i.e., wake, Stage 1, Stage 2, SWS, and REM) were reclassified in continuative and undisturbed 3-min bins and the bins were used for further analysis.

#### - Data reduction

Ten subjects were excluded from further analyses for the following reasons: 1) Four subjects did not have ECG recordings, 2) three subjects had disconnected/loose ECG reference electrodes and we were not able to detect heart beats from those subjects, 3) Three subjects had no 3-min Stage 2 and SWS epochs. Nine subjects out of the 45 subjects did not have any 3-min continuous epochs of SWS. This led to lower number of samples for SWS group. Two subjects from Stage 2 group were also considered as outliers, due to artifacts. The average number of 3-min bins for Wake, Stage 2, SWS, and REM were 4.95, 9.34, 6.11, and 3.27, respectively.

#### - Heart-beat detection and time-series extraction

The ECG signals were filtered with a passband of 0.5-100 Hz by Butterworth filter. R waves were identified in the ECG using the Pan-Tompkins method ^51^, and confirmed with visual inspection. In order to extract continuous RR tachograms, the RR intervals were resampled (at 4 Hz for power spectrum estimation; at 256 Hz for co-modulogram analysis) and interpolated by piecewise cubic spline. Zero-phase Butterworth filters were applied to the interpolated RR time-series to extract RR_LF_ and RR_HF_.

#### - HR burst detection

Within 3-min bins during wake and sleep stages, the HR burst events were detected as the minima in RR time-series with amplitude greater than two standard deviations below the mean of the RR time-series.

#### - Power spectra

The EEG power spectrum was computed using the Welch method (4 sec Hanning windows with 50 % overlap) 52. For RR time-series, the power spectral estimation was performed by the autoregressive model and the model order was set at 16 ^53^.

### C. Phase-amplitude analysis

For a given frequency pair in each stage, the RR time-series (slow or phase-providing signal) and the EEG signal (fast signal or amplitude-providing frequency) were filtered (zero-phase infinite-impulse-response bandpass filters). Phase-providing frequencies ranged from 0.04-.4 Hz (0.01 Hz increments, 0.02 Hz filter bandwidth) and amplitude-providing frequencies ranged from 0.25-16 Hz (0.5 Hz increments, 1 Hz filter bandwidth). The Hilbert transform was applied to the 3-min binned data. EEG amplitude and RR phase were then extracted and concatenated across the bins to construct the amplitude and phase time series, respectively. The phase time-series were binned into 36 10º bins (*nbins*=36) and the mean of the EEG amplitude over each bin was calculated and then normalized by dividing it by the sum over the bins. Given the normalized amplitude distribution, *P*, the modulation index (MI) was calculated by dividing the Kullback–Leibler distance ^26^ of distribution P from the uniform distribution (*U*) by log (*nbins*). We then computed for each frequency pair the normalized modulation index (nMI) by generating surrogate MIs based on the method provided in supplement of ^54^.

For statistical analysis, the nMIs were recalculated in 8 frequency pairs: LF (0.04–0.15 Hz)–delta (0.05–4 Hz), LF–theta (4–8 Hz), LF–alpha (8–13 Hz), LF–sigma (12–15 Hz), HF (0.15–0.4 Hz)–delta, HF–theta, HF–alpha, and HF–sigma.

The preferred phase for LF–delta modulation was calculated as angle of the average composite signal of A_Delta_(t) exp(i ϕ_LF_(t)) ^54^.

### D. Event-based analysis

#### - Slow oscillation

The EEG signals of frontal and central channels were filtered (zero-phase bandpass, .1–4 Hz). Then, the SO were detected based on a set of criteria for peak-to-peak amplitude, up-state amplitude, and duration of down- and up-states ^55^.

#### - Spindles

Sleep spindles were detected by applying the wavelet transform, using an 8-parameter complex Morlet wavelet with center frequency 13.5 Hz and calculating the moving average in 100 ms sliding windows. A spindle event was identified whenever the rectified signal exceeded threshold ^56^.

#### - SO-spindle coupling

The SO-spindle coupling was measured by calculation of modulation index ^26^ between spindle activity (Morlet wavelet-filtered sigma activity ^56^) and filtered (0.1–4 Hz) SO phase (within 2.5-sec windows centered at the SO negative peak).

#### - Time-locked analysis

In order to calculate changes in delta and sigma power around the HR burst, the Hilbert transform was applied on filtered EEG signals in bands of interest (0.5–4 Hz for SWA and 12–15 Hz for sigma band). To assess the HF amplitude change around the HR burst, the Hilbert transform was applied on RR_HF_.

#### - Event-locked averaging

The analyses were subject-based. We first performed a within-subject average and then averaged those averages across subjects.

#### -Change scores

The density of SOs (number of SOs per minute; SO negative peak was considered for counting) and spindles (spindle maximum amplitude point was considered for counting), as well as average delta and sigma amplitudes were calculated in both the 5-sec window prior the HR bursts, as modulated values, and in periods with no HR burst, as reference (baseline) values. That is, the baseline activities were calculated by excluding the 20-sec periods (the segments around the HR bursts) from the entire stage data. For example, if we found 6 HR bursts during 9 minutes of Stage 2, the baseline Stage 2 SWA would be calculated over 7 minute of non-bursting periods (9-6*20s/60s). For HF change score, the modulated values were calculated in the 5-sec window after the HR bursts. The subtraction between modulated and reference values divided by summation of those values were calculated as the changes scores.

### E. Statistics summary

Statistical analyses were conducted using MATLAB 2015b (MathWorks). p<.05 was considered significant.

#### - Controlling the false discovery rate

In order to control for multiple comparisons, we implemented the Benjamini-Hochberg procedure of false discovery rate (FDR) correction with q = 0.05 ^57^.

#### -Comodulogram analysis

For wake and sleep stage, the nMI was calculated in 8 frequency pairs: LF–SWA, LF-theta, LF–alpha, LF–sigma, HF–SWA, HF–theta, HF–alpha, and HF–sigma. Within each sleep stage, the nMI pairs were compared using one-way ANOVA. The Kruskal-Wallis test was used to compare a specific nMI across sleep stages. Two-tailed unpaired-samples t-test was used to compare nMI of a specific frequency pair across two sleep stage.

#### - Event-locked analysis

For wake and sleep stages, repeated measures ANOVAs followed by post hoc paired-samples t-tests were used to compare SWA as well as sigma activity across non-bursting periods and four 5-sec bins around the HR bursts.

#### - Test of periodicity of HR bursts

We calculated the inter-HRB-intervals (IHBI). For a periodic system, the Poincaré map (i.e., return map) of IHBI_n+1_ versus IHBI_n_ contains one point. In a corresponding biological system with noise, this single point becomes a normally distributed single-point cluster. The Poincaré map data was considered to be a normally distributed single-point cluster if the data points along the Poincaré map axes passed a test for normality ^57^.

### F. Memory performance tests

#### -Face-Name association (FNA) task

Face stimuli were chosen from a UC Riverside IRB-approved database of photographs of highly diverse UC Riverside undergraduate students. All students whose photographs were included in the database provided informed consent for their picture to be used to create experimental stimuli. All faces were forward-facing, shown from the shoulders up against a plain gray background, and edited to be gray scale. First and last names were selected from the 2010 United States Census data. The five most frequent male names, female names, and last names (e.g., Smith) were eliminated. Unisex names that are commonly used for both men and women were also eliminated, as were last names that are commonly used as first names (e.g., Thomas) or contain a common first name base (e.g., *Richard*son). Individual face-name pairings were randomly generated for each participant so that no two participants saw the same face-name pairs. During session 1, 44 faces (22 men and 22 women) were presented in the center of the screen with a first and last name shown below the face. The first two and last two face-name pairs were discarded after encoding (i.e., not tested) due to primacy and recency effects. Each face/name pair was presented four times for duration of 4000ms, with an inter-stimulus-interval of 500ms. Subjects were instructed to view each face-name pair and to do their best to remember each person’s name for a later test. Immediately following encoding, as well as after an 8-hr retention interval, subjects completed a recall memory test. During recall, 10 faces were presented and subjects were asked to recall the first and last names associated with each face. That is, they had the opportunity to recall 20 names (first and last names) total in each session. The ten faces tested in session 2 were different than those tested in session 1, the remaining 20 faces were not used. Total recall was calculated as the sum of first and last names recalled correctly. The difference in total recall between sessions (PM-AM) was calculated as a measure of declarative memory consolidation.

#### -Texture discrimination task (TDT)

Subjects performed a texture discrimination task similar to that developed by Karni & Sagi ^59^. Visual stimuli for the TDT were created using the Psychophysics Toolbox ^60^. Each stimulus contained two targets: a central letter (‘T’ or ‘L’), and a peripheral line array (vertical or horizontal orientation) in one of four quadrants (lower left, lower right, upper left, or upper right) at 2.5°–5.9° eccentricity from the center of the screen. The quadrant was counterbalanced across subjects. The peripheral array consisted of three diagonal bars that were either arranged in a horizontal or vertical array against a background of horizontally oriented background distracters, which created a texture difference between the target and the background.

An experimental *trial* consisted of the following sequence of four screens: central fixation cross, target screen for 33 ms, blank screen for a duration between 17 and 600 ms (the inter-stimulus-interval, or ISI), mask for 17 ms, followed by the response time interval (2000 ms) and feedback (250 ms, red fixation cross with auditory beep for incorrect trials and green fixation cross for correct trials) before the next trial. Subjects discriminated two targets per trial by reporting both the letter at central fixation (‘T’ or ‘L’) and the orientation of the peripheral array of three diagonal lines (horizontal or vertical) by making two key presses. The central task controlled for eye movements.

Each *block* consisted of 25 trials, each with the same ISI. A threshold was determined from the performance across 13 blocks, with a progressively shorter ISI, starting with 600ms and ending with 0 ms. The specific sequence of ISIs across an entire session was (600, 500, 400, 300, 250, 200, 167, 150, 133, 100, 67, 33, 17). A psychometric function of percent correct for each block was fit with a Weibull function to determine the ISI at which performance yielded 80% accuracy. TDT performance was calculated as the difference in threshold between Session 1 and Session 2, such that a positive score indicates performance improvement (i.e., decreased threshold in Session 2), whereas a negative score indicates deterioration ^61^.

Subjects were given task instructions and practiced the task during an orientation appointment prior to starting the study. During this practice, the peripheral target was located in a quadrant that was not used during the study. This practice ensured that subjects understood the task and accomplished the general task learning that typically occurs the first time a subject performs a task. Additionally, on the study day, subjects were allowed to practice an easy version of the task (ISI of 1000-600ms) prior to starting the test session to make sure subjects were able to discriminate the peripheral target between 90% and 100% correct on an easy version of the task.

## Acknowledgement

This work was supported by grants from NIH (R01AG046646), ONR (MURI: N000141310672), and Young Investigator Prize to Sara C. Mednick. MN would like to thank Nicola Cellini and Lauren N. Whitehurst for their skillful help in statistical analysis and data interpretation. Maryam Ahmadi, Negin Sattari, Benjamin D Yetton are thanked for reading through drafts of the manuscript. The authors declare no conflict of interest.

## Author Contributions

Conceived and designed the experiments: SCM EAM. Performed the experiments: SCM EAM. Analyzed the data: MN GPK EAM. Wrote the paper: MN GPK MB SCM.

## Supplementary Information

### i.) Test of periodicity of HR bursts

We tested the periodicity of the HR bursts by calculating the inter-HRB-intervals (IHBI). For a periodic system, the Poincaré map (i.e., return map) of IHBI_n+1_ versus IHBI_n_ contains one point. In a corresponding biological system with noise, this single point becomes a normally distributed single-point cluster. The form of our Poincare´ map was considered to be a normally distributed single-point cluster if the datapoints along the Poincaré map axes passed a test for normality. Shapiro-Wilk test showed 38 out of 45 subjects did not have periodic HR bursts during Stage 2 (Figure S1a). The Chi-square test showed 43 out of 45 subjects did not have periodic HR bursts during Stage 2. In Summary, our conclusion is that the HR bursts are less likely to be periodic events.

### ii.) Exploratory analyses on theta and beta activity

Repeated measures ANOVAs indicated significant differences in theta activity across the 5-sec bins and periods with no HR burst only during Stage 2 (F_4,168_=52.36, p<.00001, Supp Fig. 4; it was also shown in comodulogrms in Fig 3e). The highest theta power occurred during the 5-sec bin prior the HR burst (t(84)=5.40, p<.00001). Repeated measures ANOVAs also indicated significant differences in beta activity across the 5-sec bins and periods with no HR burst during Wake (F_4,132_=12.82, p<.00001, Supp Fig. 4) and Stage 2 (F_4,168_=70.48, p<.00001). Post-hoc comparisons revealed the highest beta power occurred in the 5-sec bin prior the HR burst during Wake (t(66)=3.58, p=.0006) as well as during Stage 2 (t(84)=5.35, p<.00001).

In contrast with SWA and Sigma power, no significant correlation was found between the recall difference score and ACE-related increases in theta power (r=0.061, p=.697) and beta power (r=0.038, p=.812) during Stage 2. In addition, no significant correlation was found between the recall difference score and ACE-related increases in beta power during Wake (r=-0.230, p=.190).

### iii.) Exploratory analyses on the application of modulation index between HR bursts and EEG activities rather than the EEG change scores

We calculated the modulation index (MI; Tort et al 2008) between the phase of heart rate bursts and amplitude of slow wave activity and sigma power within 20-sec windows around the HRBs during Stage 2 and SWS. The recall difference score in the declarative memory task was positively correlated with HRB-SWA MI in Stage 2 (r= .339, p= .0246) and SWS (r= .334, p= .0468), but non-significant with HRB-Sigma power MI in Stage 2 (r=0.248, p=0.1050). The correlation between the recall difference score and HRB-Sigma power MI in SWS was not significant (r= -0.002, p=.990).

For the procedural learning task, the discrimination improvement was negatively correlated with HRB-SWA MI in Stage 2 (r= -.3965, p= .0077) and, marginally, SWS (r=-0.3064, p=0.0691). The correlations between the discrimination improvement score and HRB-Sigma power MI in Stage 2 (r= -.0294, p= .8514) and SWS (r=-.0588, p=0.7371) were not significant

**Supplemental Figure 6.**
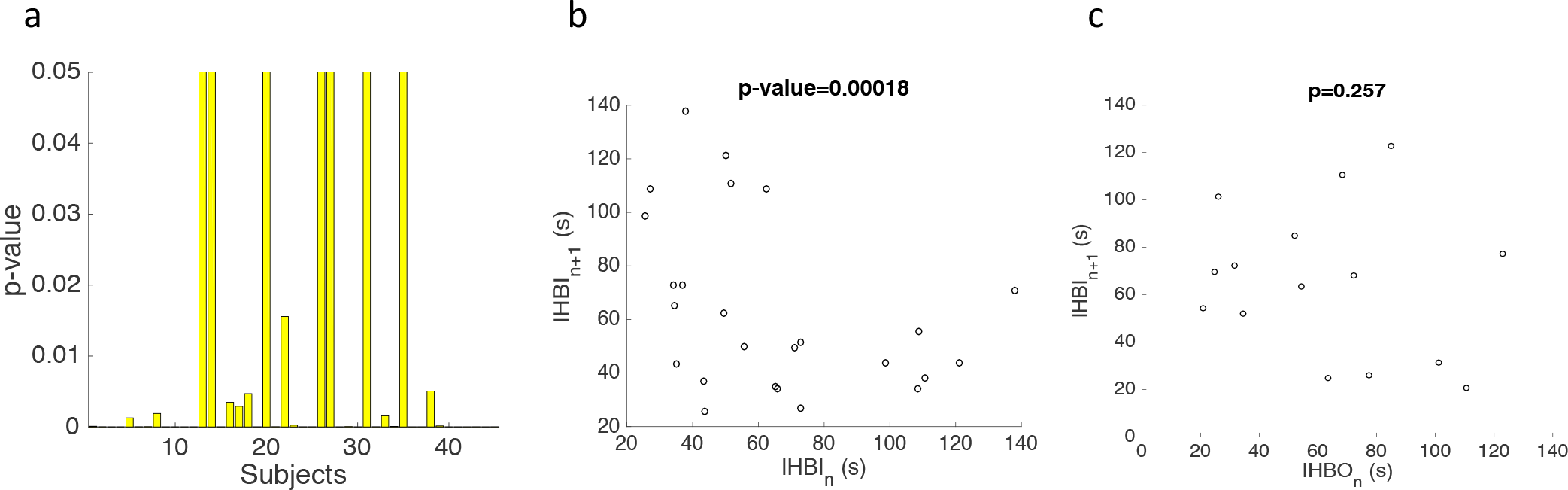
Test of periodicity of HR bursts during Stage 2. a) Shapiro-Wilk test shows only 7 subjects out of 45 subjects do not reject the null hypothesis of HRB periodicity. b) Poincaré map for aperiodic inter-HRB-intervals in one subject. c) Poincaré map for periodic inter-HRB-intervals in one subject.

**Supplemental Figure 7.**
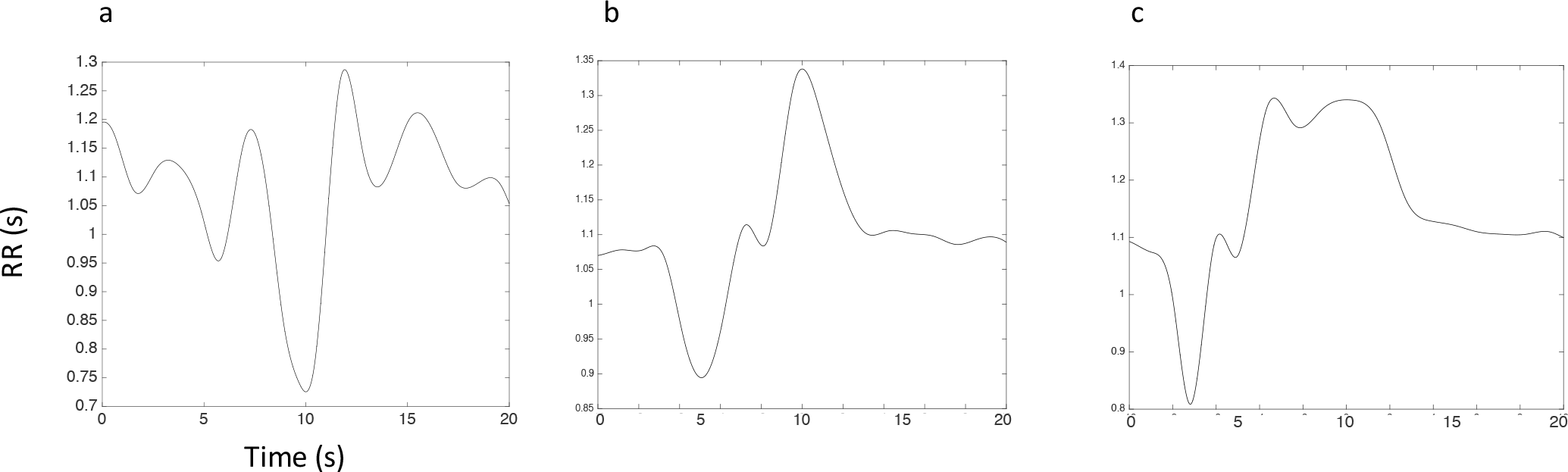
An example of three kinds of HR bursts in one subject. HR decelerations after HR bursts can be a) small to detect and b-c) change in duration.

**Supplemental Figure 8.**
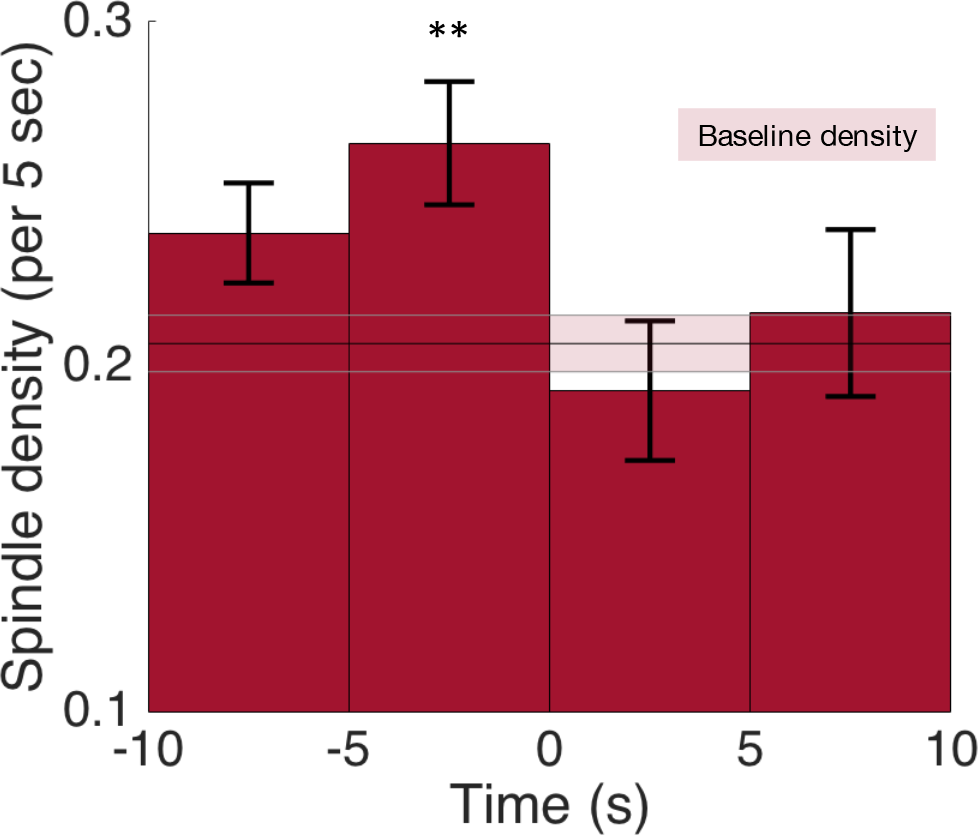
The density of sleep spindles increases in the 5-sec bin prior to the peak of HR bursts during Stage 2.

**Supplemental Figure 9.**
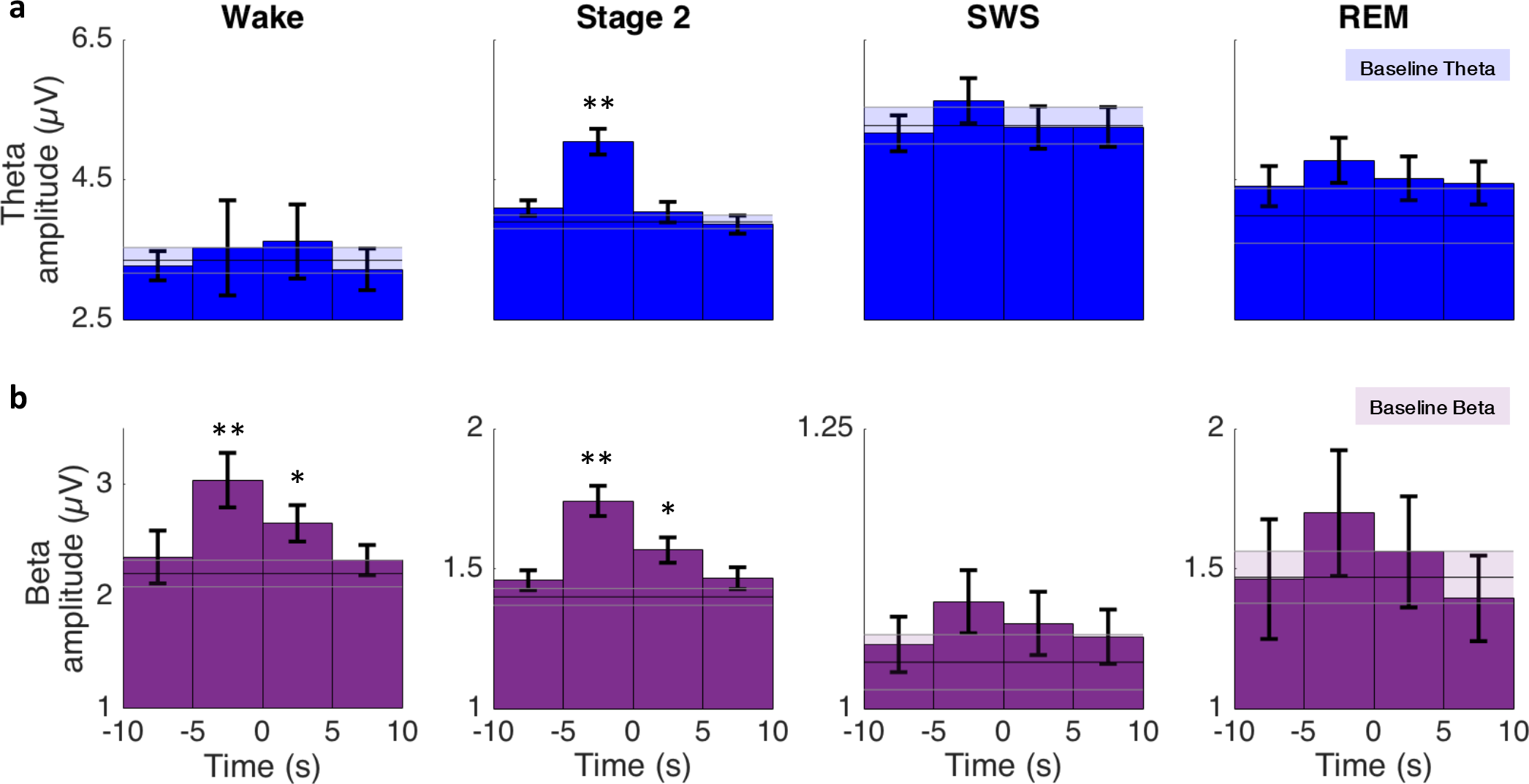
The event-related analysis of changes in a) theta and b) beta activities around HR bursts. Asterisks in show the significant differences after FDR correction (*p<.05 and **p<.001).

**Supplemental Figure 10.**
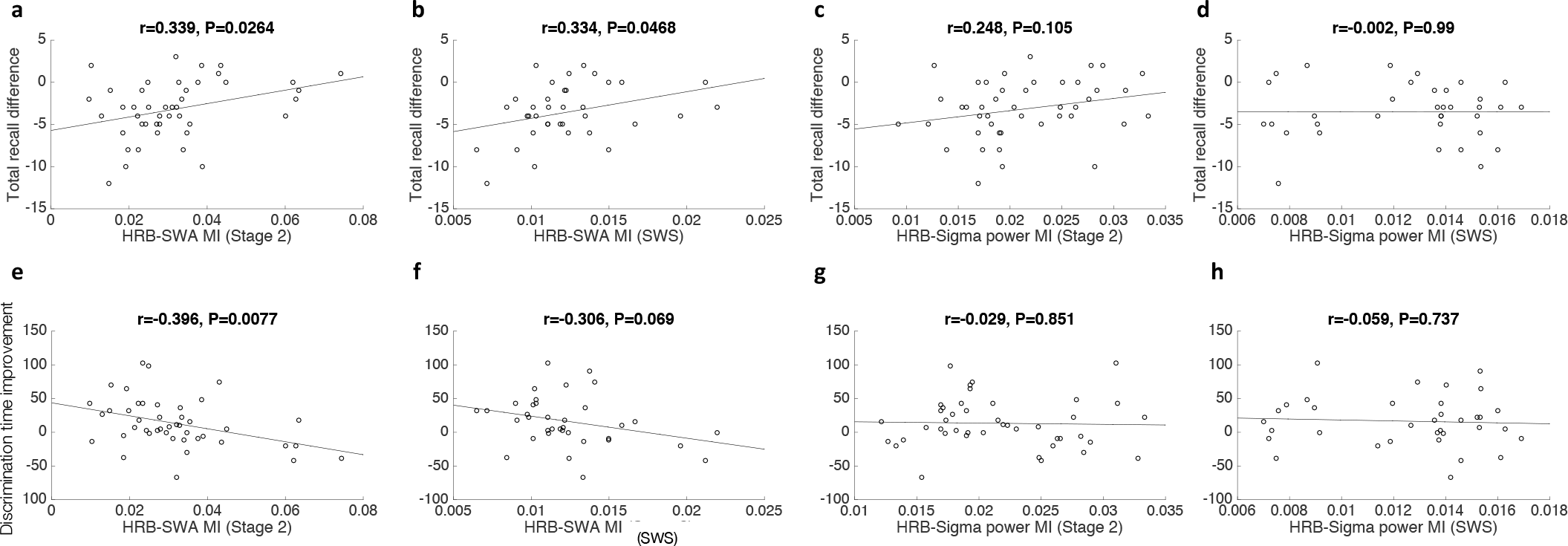
a-d) Scatter plots for relationships between the recall improvement in the declarative memory (face-name task) and HRB-SWA/sigma modulation index during Stage 2 and SWS. e-h) Scatter plots for relationships between improvement in the perceptual learning (texture discrimination task) and HRB-SWA/sigma modulation index during Stage 2 and SWS.

## References

1. Gais, S. & Born, J., Declarative memory consolidation: Mechanisms acting during human sleep. Learning and Memory 11, 679-685 (2004).

2. Girardeau, G. & Zugaro, M., Hippocampal ripples and memory consolidation. Current Opinion in Neurobiology 21 (3), 452-459 (2011).

3. Nir, Y. et al., Regional Slow Waves and Spindles in Human Sleep. Neuron 70 (1), 153–169 (2011).

4. Mölle, M., Bergmann, T. O., Marshall, L. & Born, J., Fast and Slow Spindles during the Sleep Slow Oscillation: Disparate Coalescence and Engagement in Memory Processing. Sleep 34 (10), 1411–1421 (2011).

5. Mölle, M., Eschenko, O., Gais, S., Sara, S. J. & Born, J., The influence of learning on sleep slow oscillations and associated spindles and ripples in humans and rats. European Journal of Neuroscience 29 (5), 1071-1081 (2009).

6. Niknazar, M., Krishnan, G. P., Bazhenov, M. & Mednick, S. C., Coupling of Thalamocortical Sleep Oscillations Are Important for Memory Consolidation in Humans. Plos One 10 (12), e0144720 (2015).

7. Mednick, S. C. et al., The critical role of sleep spindles in hippocampal-dependent memory: a pharmacology study. The Journal of Neuroscience 33 (10), 4494-4504 (2013).

8. Staresina, B. P. et al., Hierarchical nesting of slow oscillations, spindles and ripples in the human hippocampus during sleep. Nature neuroscience 18, 1679–1686 (2015).

9. McGaugh, J. L., Making lasting memories: remembering the significant. Proc Natl Acad Sci U S A 110(Suppl 2), 10402-10407 (2013).

10. Packard, M. G., Williams, C. L., Cahill, L. & McGaugh, J. L., in Neurobehavioral Plasticity: Learning, Development and Response to Brain Insults (Lawrence Erlbaum Associates;, New Jersey, 1995), pp. 149–184.

11. Thayer, J. F. & Lane, R. D., Claude Bernard and the heart-brain connection: Further elaboration of a model of neurovisceral integration. Neuroscience and Biobehavioral Reviews 33 (2), 81-88 (2009).

12. Williams, C. L. & Jensen, R. A., Effects of vagotomy on Leu-enkephalin-induced changes in memory storage processes. Physiol Behav 54 (4), 650-663 (1993).

13. Clark, K. B., Naritoku, D. K., Smith, D. C., Browning, R. A. & Jensen, R. A., Enhanced recognition memory following vagus nerve stimulation in human subjects. Nat Neurosci 2 (1), 94-98 (1999).

14. Whitehurst, L. N., Cellini, N., McDevitt, E. A., Duggan, K. A. & Mednick, S. C., Autonomic activity during sleep predicts memory consolidation in humans. Proceedings of national academy of science 113 (26), 7272–7277 (2016).

15. de Zambotti, M. et al., K-Complexes: Interaction between the Central and Autonomic Nervous Systems during Sleep. Sleep 39 (5), 1129–1137 (2016).

16. Fruhistorfer, H., Partanen, J. & Lumio, J., Vertex sharp waves and heart action during the onset of sleep. Electroencephalogr Clin Neurophysiol 31 (6), 614-617 (1971).

17. Lechinger, J., Heib, D. P., Gruber, W., Schabus, M. & Klimesch, W., Heartbeat-related EEG amplitude and phase modulations from wakefulness to deep sleep: Interactions with sleep spindles and slow oscillations. Psychophysiology 52 (11), 1441-1450 (2015).

18. Gray, M. A. et al., A cortical potential reflecting cardiac function. Proc Natl Acad Sci U S A 104 (16), 6818-6823 (2007).

19. Ako, M. et al., Correlation between electroencephalography and heart rate variability during sleep. Psychiatry and Clinical Neurosciences 57 (1), 59-65 (2003).

20. Norton, K. N., Luchyshyn, T. A. & Shoemaker, J. K., Evidence for a medial prefrontal cortex–hippocampal axis associated with heart rate control in conscious humans. Brain Research, 104-105 (2013).

21. Pedemonte, M., Goldstein-Daruech, N. & Velluti, R. A., Temporal correlations between heart rate, medullary units and hippocampal theta rhythm in anesthetized, sleeping and awake guinea pigs. Autonomic Neuroscience: Basic and Clinical 107 (2), 99-104 (2003).

22. Malik, M. et al., Heart rate variability. Standards of measurement, physiological interpretation, and clinical use. European Heart Journal 17 (3), 354-381 (1996).

23. Billman, G. E., The LF/HF ratio does not accurately measure cardiac sympatho-vagal balance. Frontiers in Physiology 4, 1-5 (2013).

24. Vanoli, E. et al., Heart rate variability during specific sleep stages. A comparison of healthy subjects with patients after myocardial infarction. Circulation 91 (7), 1918-1922 (1995).

25. Cellini, N., Whitehurst, N. L., McDevitt, E. A. & Mednick, S. C., Heart rate variability during daytime naps in healthy adults: Autonomic profile and short-term reliability. Psychophysiology 53 (4), 473–481 (2016).

26. Tort, A. B., Komorowski, R., Eichenbaum, H. & Kopell, N., Measuring Phase-Amplitude Coupling Between Neuronal Oscillations of Different Frequencies. Journal of neurophysiology 104 (2), 1195-1210 (2010).

27. Karni, A., Tanne, D., Rubenstein, B. S., Askenasy, J. J. & Sagi, D., Dependence on REM sleep of overnight improvement of a perceptual skill. Science 265 (5172), 679-682 (1994).

28. Rothenberger, S. D. et al., Time-varying correlations between delta EEG power and heart rate variability in midlife women: the SWAN Sleep Study. Psychophysiology 52 (4), 572-584 (2015).

29. Brandenberger, G., Ehrhart, J., Piquard, F. & Simon, C., Inverse coupling between ultradian oscillations in delta wave activity and heart rate variability during sleep. Clinical Neurophysiology 112 (6), 992-996 (2001).

30. Mensen, A., Zhang, Z., Qi, M. & Khatami, R., The occurrence of individual slow waves in sleep is predicted by heart rate. Scientific Reports 6, 29671 (2016).

31. Rowe, K. et al., Heart rate surges during REM sleep are associated with theta rhythm and PGO activity in cats. American Journal of Physiology 277 (3), R843-R849 (1999).

32. Datta, S., Activation of phasic pontine-wave generator: A mechanism for sleep-dependent memory processing. Sleep and Biological Rhythms 4, 16-26 (2006).

33. Terzano, M. G. et al., The Cyclic Alternating Pattern as a physiologic component of normal NREM sleep. Sleep 8 (2), 137-145 (1985).

34. Lecci, S. et al., Coordinated infraslow neural and cardiac oscillations mark fragility and offline periods in mammalian sleep. Science Advances 3 (e1602026), 1-14 (2017).

35. Chapleau, M. W. & Sabharwal, R., Methods of assessing vagus nerve activity and reflexes. Heart Fail Rev 16 (2), 109-127 (2011).

36. Guyenet, G., The sympathetic control of blood pressure. Nature Reviews Neuroscience 7, 335-346 (2006).

37. Barraco, I. R. A., Nucleus of the Solitary Tract (CRC Press, 1993).

38. van der Kooy, D., Koda, L. Y., McGinty, J. F., Gerfen, C. R. & Bloom, F. E., The Organization of Projections From the Cortex, Amygdala, and Hypothalamus to the Nucleus of the Solitary Tract in Rat. The Journal of Comparative Neurology 224 (1), 1-24 (1994).

39. Samuels, E. R. & Szabadi, E., Functional neuroanatomy of the noradrenergic locus coeruleus: its roles in the regulation of arousal and autonomic function part I: principles of functional organisation. Curr Neuropharmacol 6 (3), 235-253 (2008).

40. Martínez-Vargas, D., Valdés-Cruz, A., Magdaleno-Madrigal, V. M., Fernández-Mas, R. & Almazán-Alvarado, S., Effect of Electrical Stimulation of the Nucleus of the Solitary Tract on Electroencephalographic Spectral Power and the Sleep–Wake Cycle in Freely Moving Cats. Brain Stimulation. Brain Stimulation 10 (1), 116-125 (2016).

41. Krishnan, G. P. et al., Cellular and neurochemical basis of sleep stages in the thalamocortical network. eLife 5 (e18607) (2016).

42. Roozendaal, B. & McGaugh, J. L., Memory Modulation. Behav Neurosci 125 (6), 797-824 (2012).

43. Engineer, N. D. et al., Reversing pathological neural activity using targeted plasticity. Nature 470 (7332), 101–104 (2011).

44. Engineer, C. T., Engineer, N. D., Riley, J. R., Seale, J. D. & Kilgard, M. P., Pairing Speech Sounds With Vagus Nerve Stimulation Drives Stimulus-specific Cortical Plasticity. Brain Stimul 8 (3), 637-644 (2015).

45. Peña, D. F., Engineer, N. D. & McIntyre, C. K., Rapid remission of conditioned fear expression with extinction training paired with vagus nerve stimulation. Biological Psychiatry 73 (11), 1071–1077 (2013).

46. Sun, L. et al., Vagus nerve stimulation improves working memory performance. Journal of Clinical and Experimental Neuropsychology, 1-11 (2017).

47. Sattari, N. et al., The effect of sex and menstrual phase on memory formation during a nap. Neurobiology of Learning and Memory 145, 119-128 (2017).

48. Johns, M. W., Daytime sleepiness, snoring, and obstructive sleep apnea. The Epworth Sleepiness Scale. Chest 103 (1), 30-36 (1993).

49. Adan, A. & Almirall, H., Horne and östberg morningness-eveningness questionnaire: a reduced scale. Person Individ Diff 12 (3), 241-253 (1991).

50. Rechtschaffen, A. & Kales, A., A manual of standardized terminology, techniques and scoring system for sleep stages of human subjects (Brain Information Services/Brain Research Institue, University of California, Los Angeles, 1968).

51. Pan, J. & Tompkins, W. J., A real-time QRS detection algorithm. IEEE Trans Biomed Eng 32 (3), 230-236 (1985).

52. Campbell, I. G., EEG Recording and Analysis for Sleep Research. Curr Protoc Neurosci 49 (10.2), 10.2.1–10.2.19 (2009).

53. Boardman, A., Schlindwein, F. S., Rocha, A. P. & Leite, A., A study on the optimum order of autoregressive models for heart rate variability. Physiol Meas 23 (2), 325-336 (2002).

54. Canolty, R. T. et al., High gamma power is phase-locked to theta oscillations in human neocortex. Science 313 (5793), 1626-1628 (2006).

55. Dang-Vu, T. T. et al., Spontaneous neural activity during human slow wave sleep. Proceeding of national academy of sciences 105 (39), 15160-15165 (2008).

56. Wamsley, E. J. et al., Reduced Sleep Spindles and Spindle Coherence in Schizophrenia: Mechanisms of Impaired Memory Consolidation? Biol Psychiatry 71 (2), 154-161 (2012).

57. Del Negro, C. A., Wilson, C. G., Butera, R. J., Rigatto, H. & Smith, J. C., Periodicity, Mixed-Mode Oscillations, and Quasiperiodicity in a Rhythm-Generating Neural Network. Biophys J 82 (1 Pt 1), 206-214 (2002).

58. Benjamini, Y. & Hochberg, Y., Controlling the False Discovery Rate: A Practical and Powerful Approach to Multiple Testing. J R Statist Soc B 57 (1), 289-300 (1995).

59. Karni, A. & Sagi, D., Where practice makes perfect in texture discrimination: Evidence for primary visual cortex plasticity. Proceedings of the National Academy of Sciences USA 88, 4966-4970 (1991).

60. Brainard, D. H., The Psychophysics Toolbox. Spatial Vision 10, 433-436 (1997).

61. Mednick, S. C., Arman, A. C. & Boynton, G. M., The time course and specificity of perceptual deterioration. Proceedings of the National Academy of Sciences USA 102 (10), 3881-3885 (2005).

